# Delineation of complex gene expression patterns in single cell RNA-seq data with ICARUS v2.0

**DOI:** 10.1101/2023.01.23.525100

**Authors:** Andrew Jiang, Linya You, Russell G Snell, Klaus Lehnert

## Abstract

Complex biological traits and disease often involve patterns of gene expression that can be characterised and examined. Here we present ICARUS v2.0, an update to our single cell RNA-seq analysis web server with additional tools to investigate gene networks and understand core patterns of gene regulation in relation to biological traits. ICARUS v2.0 enables gene co-expression analysis with MEGENA, transcription factor regulated network identification with SCENIC, trajectory analysis with Monocle3, and characterisation of cell-cell communication with CellChat. Cell cluster gene expression profiles may be examined against Genome Wide Association Studies with MAGMA to find significant associations with GWAS traits. Additionally, differentially expressed genes may be compared against the Drug-Gene Interaction database (DGIdb 4.0) to facilitate drug discovery. ICARUS v2.0 offers a comprehensive toolbox of the latest single cell RNA-seq analysis methodologies packed into an efficient, user friendly, tutorial style web server application (accessible at https://launch.icarus-scrnaseq.cloud.edu.au/) that enables single cell RNA-seq analysis tailored to the user’s dataset.

## Introduction

Many biological networks contain patterns of gene expression that are evolutionary conserved. These patterns of networks are comprised of intertwined signalling pathways that are fundamentally important for biological functionality including development, differentiation, response to stimuli and senescence. During disease, many of these pathways become dysregulated and characterisation of these affected cell populations and their regulatory networks is vital for understanding disease mechanisms and pathogenesis (1–3). Here, we present an update to ICARUS (4), our web server tool to enable users without experience in R to undertake single cell RNA-seq analysis and aid hypothesis generation. ICARUS v2.0 assembles the latest single cell RNA-seq gene network construction and analysis tools to allow interpretation and deconstruction of complex biological traits.

ICARUS v2.0 supports establishment of gene networks through the implementation of; Multiscale Embedded Gene Co-expression Network Analysis (MEGENA) (5), transcription factor regulatory network construction through single-cell regulatory network inference and clustering (SCENIC) (6), characterisation of gene expression changes through pseudotime (cell trajectories) with Monocle3 (7–9) and cell-cell communication signalling classification through identification of ligand-receptor pairs with CellChat (10). Furthermore, the gene expression profiles of cell populations can be examined for associations with Genome Wide Association Study (GWAS) traits using the multi-marker analysis of genomic annotation (MAGMA) tool (11,12) to explore crucial cell populations that may drive the affected phenotype. Finally, differentially expressed genes can be queried against the drug-gene interactions database, DGIdb 4.0 (13) to facilitate the identification of drug targets for therapeutic testing.

Further updates to data processing methods are also included in this update. These include, a quality control step to identify cell doublets/multiplets using DoubletFinder (14), an improved method of data normalisation, SCTransform (15,16), and a new method for cell cluster labelling, sctype (17) that congregates cell type specific markers from CellMarker (18) and PanglaoDB (19) databases. Summary of R packages used in ICARUS v2.0 including the main command are detailed in Table 1. ICARUS v2.0 is accessible through an efficient, userfriendly web server at https://launch.icarus-scrnaseq.cloud.edu.au/.

**Table 1.**
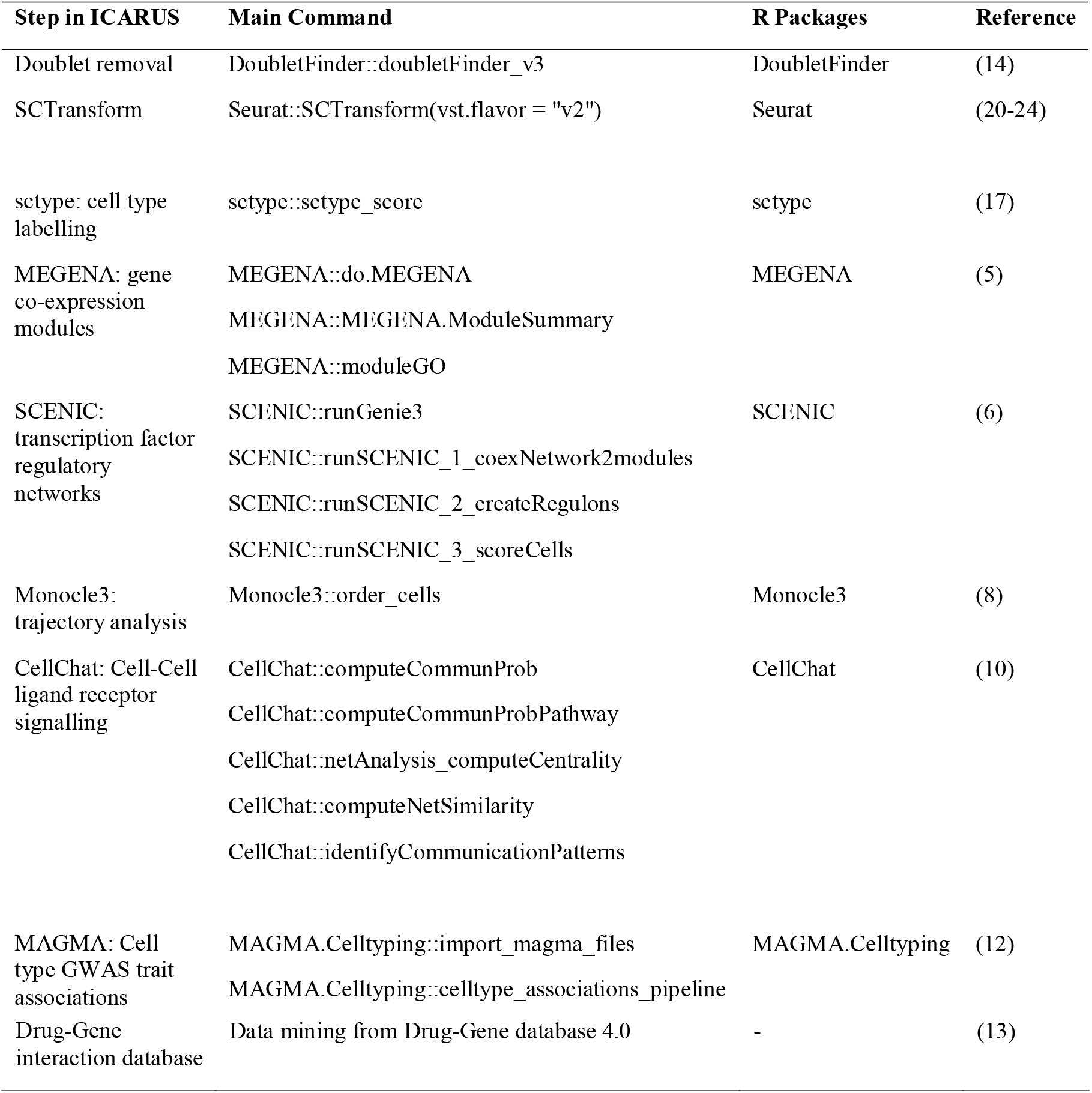
Summary of R packages used in ICARUS v2.0.

## Materials and Methods

### Updates to processing methods

#### Doublet Removal

Cell doublets or multiplets may arise during scRNA-seq library preparation when droplet-based microfluidics methods partition multiple cells into a single droplet. These doublets will display mixed transcriptomes and can compromise downstream analysis by creating spurious intermediate populations or transition states. ICARUS v2.0 introduces the DoubletFinder R package to detect and remove heterotypic doublets (Default settings, refer to https://github.com/chris-mcginnis-ucsf/DoubletFinder) (14). DoubletFinder first simulates “artificial doublets” using transcriptional profiles from pairs of cells in real data. The artificial doublets are merged with real data and dimensionality reduction is performed. Lastly, a k-nearest neighbour graph is developed, and each cell is scored based on its proximity to artificial doublets. The highest scoring cells are assigned as real doublets. The user has the option to visualise and remove these doublets from further analysis.

#### SCTransform

Scaling and normalization of raw gene count matrices of single cell RNA-seq data is a common practice to account for technical confounders such as differences in sequence depth. ICARUS v2.0 now includes the Seurat SCTransform method of data normalisation which recovers sharper biological distinctions between formed cell clusters compared to lognormalisation. The SCTransform methodology computes Pearson residuals from a negative binomial regression of the raw count data and applies dimensionality reduction methods (15,16).

#### sctype and additional SingleR datasets

The expression profiles of marker genes for each cell cluster may be compared against existing cell marker datasets to assign cell type labels to clusters. ICARUS v2.0 introduces cell cluster labelling with sctype (17), an ultra-fast unsupervised method for cell type annotation using compiled cell markers from CellMarker (http://bio-bigdata.hrbmu.edu.cn/CellMarker/) (18) and PanglaoDB (https://panglaodb.se/) (19) databases. Sctype is computationally efficient and includes marker gene sets for 11 different tissue types including brain, pancreas, immune system, liver, eye, kidney, lung, embryonic, gastrointestinal tract, muscle and skin (17).

ICARUS provides the SingleR supervised cell-type assignment algorithm to annotate cell clusters by comparison to previously annotated cell types in single cell datasets (25). ICARUS v2.0 extends the utility of SingleR by including additional reference datasets including Darmanis human temporal lobe data (26); Zhong human embryonic prefrontal cortex data (27); Baron human and mouse pancreas data (28); Lawlor human non-diabetic and diabetic pancreas data (29); Muraro human pancreas data (30); Segerstolpe human nondiabetic and diabetic pancreas data (31) and the He organ atlas, which comprises data from 15 organs (bladder, blood, common bile duct, oesophagus, heart, liver, lymph node, bone marrow, muscle, rectum, skin, small intestine, spleen, stomach and trachea) (32).

### Gene network construction

#### MEGENA: gene co-expression modules

Patterns of biological networks can be grouped into hierarchal co-expression modules which infer functionality and dictate biological causality. A method of detecting these gene coexpression modules is multiscale embedded gene co-expression network analysis (MEGENA) which employs a planar maximally filtered graph to extract significant gene interactions and constructs co-expression gene modules (5). MEGENA first classifies gene co-expression modules through identification of significant gene interactions via planar filtered network construction and embedding on a topological sphere. Multi-scale clustering is then applied to group similar and dissimilar gene modules. Lastly, multiscale hub analysis is used to connect clusters in a hierarchical structure.

ICARUS v2.0 performs MEGENA on either a set of highly variable genes (Seurat::FindVariableFeatures) or user computed differentially expressed genes to generate a set of hierarchically ordered co-expression modules, where larger modules progressively branch into smaller submodules. A heatmap of significant Gene Ontology terms associated with each co-expression module is also provided to aid interpretation and hypothesis generation. Additionally, a hierarchical sunburst plot of computed MEGENA co-expression modules is produced which highlights the cell cluster/cell type that displays the highest activity of the corresponding module. Module activity is computed as a percentage of cluster marker gene overlap (Seurat::FindAllMarkers) with co-expression module genes.

#### SCENIC: transcription factor regulatory networks

A cell’s transcriptional state may be characterised by gene regulatory networks (GRNs) that are formed by transcription factors and cofactors that regulate each other and their downstream gene targets. ICARUS v2.0 utilises the SCENIC R package (6) to characterise cell cluster/cell type specific GRNs using either a set of highly variable genes (Seurat::FindVariableFeatures) or user computed differentially expressed genes. SCENIC performs cis-regulatory transcription factor binding motif analysis on a set of co-expressed transcription factors and variable genes. Transcription factor motifs in the promoter region of genes (up to 500bp upstream of the transcription start site) and 20kb around the transcription start site (+/- 10kb) are scored. ICARUS v2.0 currently supports SCENIC v1.2.4 which includes species specific transcription motif database for human, mouse, and fly. The activity of these transcription factor regulated gene modules is scored across cell populations using the AUCell algorithm (6). ICARUS v2.0 visualises regulated gene module activity across cell populations in 2D/3D UMAP and t-SNE plots. Heatmap and dotplot visualisations are also provided.

#### Monocle3: trajectory analysis

The expression profiles of cells change during development, in response to stimuli and throughout life. Trajectory analysis aims to determine the sequence of gene expression changes in single cell RNA-seq datasets. ICARUS v2.0 employs the Monocle3 algorithm (8) to graph cells according to their progress in pseudotime, a measure of the amount of transcriptional change a cell undergoes from its beginning to end states. Calculation of pseudotime requires the user to select a population of cells where the beginning of the biological process is likely located (known as root cells). Root cells may be selected from a specific cell population/cell type or interactively using a lasso select function. A table of genes that exhibit changes across pseudotime is also provided.

#### CellChat: Cell-Cell ligand receptor signalling

Cells interact and communicate with each other through ligand-receptor pairs that coordinate many biological processes in both healthy and diseased conditions. The characterisation of cell-cell signalling crosstalk through soluble agonists/antagonists and membrane bound cofactors may help us interpret and understand complex networks of systems biology. ICARUS v2.0 employs the CellChat (10) tool to quantitatively infer and analyse intercellular communication networks from single cell RNA-seq data. CellChat incorporates a manually curated comprehensive database of ligand-receptor pairs, soluble agonists/antagonists and stimulatory/inhibitory membrane bound co-receptors to infer cell-cell communication interactions based on social network analysis tools, pattern recognition methods and manifold learning approaches. ICARUS v2.0 currently supports CellChat v1.4.0 incorporating signalling molecule interaction databases for human and mouse. The option to analyse a single dataset or comparison between two datasets is available.

#### MAGMA: Cell type GWAS trait associations

Genome wide association studies have identified several loci and genes that are associated with a trait of interest. The expression profiles of cell populations in single cell RNA-seq data can be compared against these GWAS loci to identify potentially affected cell types underlying complex traits. ICARUS employs the multi-marker analysis of genomic annotation (MAGMA) methodology to identify increased linear association between cell population derived gene sets and gene-level GWAS summary statistics, with the hypothesis that in affected cell types, expressed genes should be more associated with the GWAS trait (11,12). The user may either upload their own GWAS summary statistics or select biological traits from over 700 public GWAS datasets from the IEU open GWAS project (https://gwas.mrcieu.ac.uk/), UK biobank (http://www.nealelab.is/uk-biobank) and FinnGen (https://www.finngen.fi/fi). Public GWAS datasets were curated by neurogenomics/MAGMA_Files_Public (https://github.com/neurogenomics/MAGMA_Files_Public) and includes gene level GWAS statistics for 10b upstream and 1.5kb downstream of each associated loci. ICARUS v2.0 utilises the MAGMA.Celltyping v.2.0.6 R package to identify cell types that may explain the heritability signal from GWAS summary statistics.

#### Drug-Gene Interaction Database

The Drug-Gene Interaction Database (DGIdb, www.dgidb.org) (13,33,34) curates information on druggable genes from publications (35,36), known affected pathways (Gene Ontology, Human Protein Atlas, IDG) and publicly available databases (DrugBank, PharmGKB, Chembl, Drug Target Commons and TTD). ICARUS v2.0 offers an option to query differentially expressed genes against DGIdb v4.0 (13) and return a list of potential drug targets with information of interaction type (i.e., inhibitor, agonist, blocker), interaction claim source (i.e., PharmGKB, ChemblInteractions, etc), interaction group score (score takes into account number of drug and gene partners and number of supporting publications) and the relevant pubmed reference if available. Suitable drugs may be tested in a laboratory setting and contribute to discovery of repurposed drugs for clinical benefit.

## Results

To demonstrate the utility of ICARUS v2.0, we have examined a single-nuclei RNA-seq data of the substantia nigra from 5 human individuals with absence of neurological clinical disease. Data is available from human cell atlas (https://data.humancellatlas.org/explore/projects/996120f9-e84f-409f-a01e-732ab58ca8b9)(37). Using this dataset, a quality control filter was first applied removing low quality cells with unique gene counts less than 200 and cells with high mitochondrial reads >5%. Dimensionality reduction with SCTransform was then performed, prioritising 3,000 highly variable genes. Substantia nigra nuclei from the 5 individuals were integrated using harmony(38) and cell clustering was performed with the first 25 dimensions, a k-nearest neighbour value of 20 and the Louvain community detection algorithm. Cluster labelling with sctype(17) identified 6 different brain cell types including oligodendrocytes (based on expression of OLIG1, OLIG2, MBP, MOG cell markers), astrocytes (based on expression of GFAP, SLC1A3, SLC1A2, AQP4 cell markers), GABAergic neurons (based on expression of GABBR1, GABBR2, GAD2, GAD1 cell markers), oligodendrocyte precursor cells (based on expression of LHFPL3, MEGF11, PCDH15, PDGFRA cell markers), endothelial cells (based on expression of CD34, EGFL7, FLT1, KDR cell markers) and microglia (based on expression of P2RY12, ITGAM, CD40, CX3CR1 cell markers) (Figure 1).

**Figure 1.**
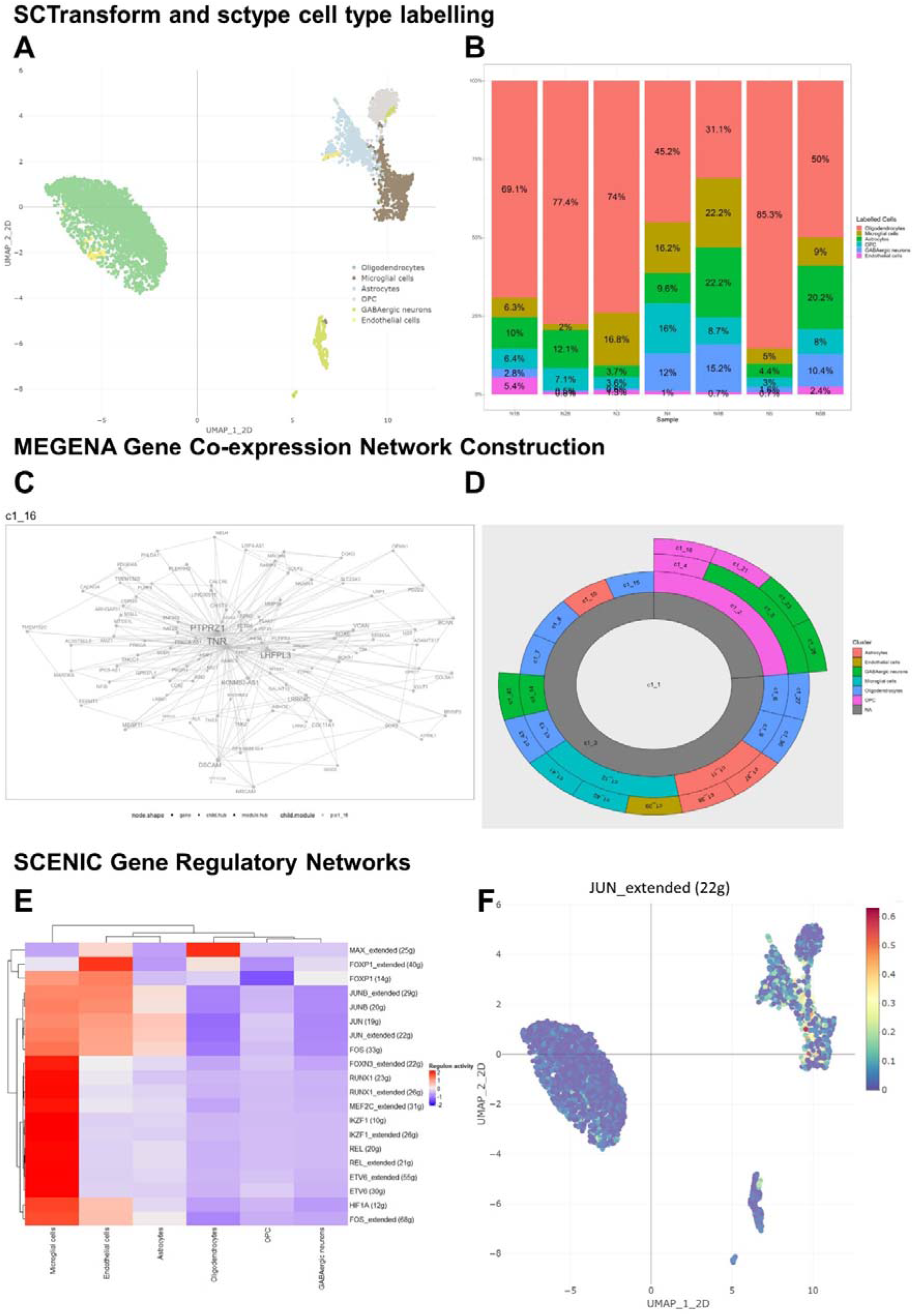
Gene network analysis of single cell RNA-seq datasets in ICARUS v2.0. Dataset analysed was from the substantia nigra of 5 healthy human individuals with an absence of neurological disease. Cell clustering with SCTransform and cell type labelling produced 7 different brain cell types (A) with their distributions shown in (B). MEGENA co-expression module construction showcased a module centred around the LHFPL3-PTPRZ1-TNR gene cluster (C) that is predominately expressed in oligodendrocyte precursor cells (D). SCENIC transcription factor regulatory network analysis (E) indicated increased JUN regulated gene module activity in the astrocytes and microglia cell clusters (F).

To showcase the gene network analysis capabilities of ICARUS v2.0, we focused on the oligodendrocyte precursor cell (OPC) cluster. MEGENA co-expression module analysis using 2,000 variable genes as input, highlighted a functional module involved in the regulation of extracellular organisation, synapse signalling and developmental morphogenesis (gene module GO analysis, supplementary figure 1) that is centred around the LHFPL3-PTPRZ1-TNR gene cluster (Figure 1C). Trajectory analysis selecting the OPCs cluster as the root cells, showcased a gradual change in pseudotime (changes in cell transcriptional state) starting from oligodendrocyte precursor cells and ending in microglia and astrocytes (Figure 2A). Top genes that change as a function of pseudotime included MEG3, DSCAM, LRRC4C (full list of genes can be found in supplementary table 1). The transition from OPC to astrocytes and microglia appears to be controlled by the cell differentiation and development related transcription factors, FOS, JUN and JUNB as determined by SCENIC transcription factor regulatory network analysis (Figure 1E). Regulated gene module activity of the JUN family of transcription factors is detailed in Figure 1F. Cell-cell communication through ligand-receptor pairs showed a greater number of interactions between OPCs and astrocytes and fewer interactions with all other cell types (Figure 2C). OPC outgoing signalling pathways included NGL, PTN, CNTN, APP and TENASCIN pathways and incoming signals from NRXN, NCAM, CNTN, PTN, NGL, PSAP and CALCR were detected (Figure 2D and Figure 2E).

**Figure 2.**
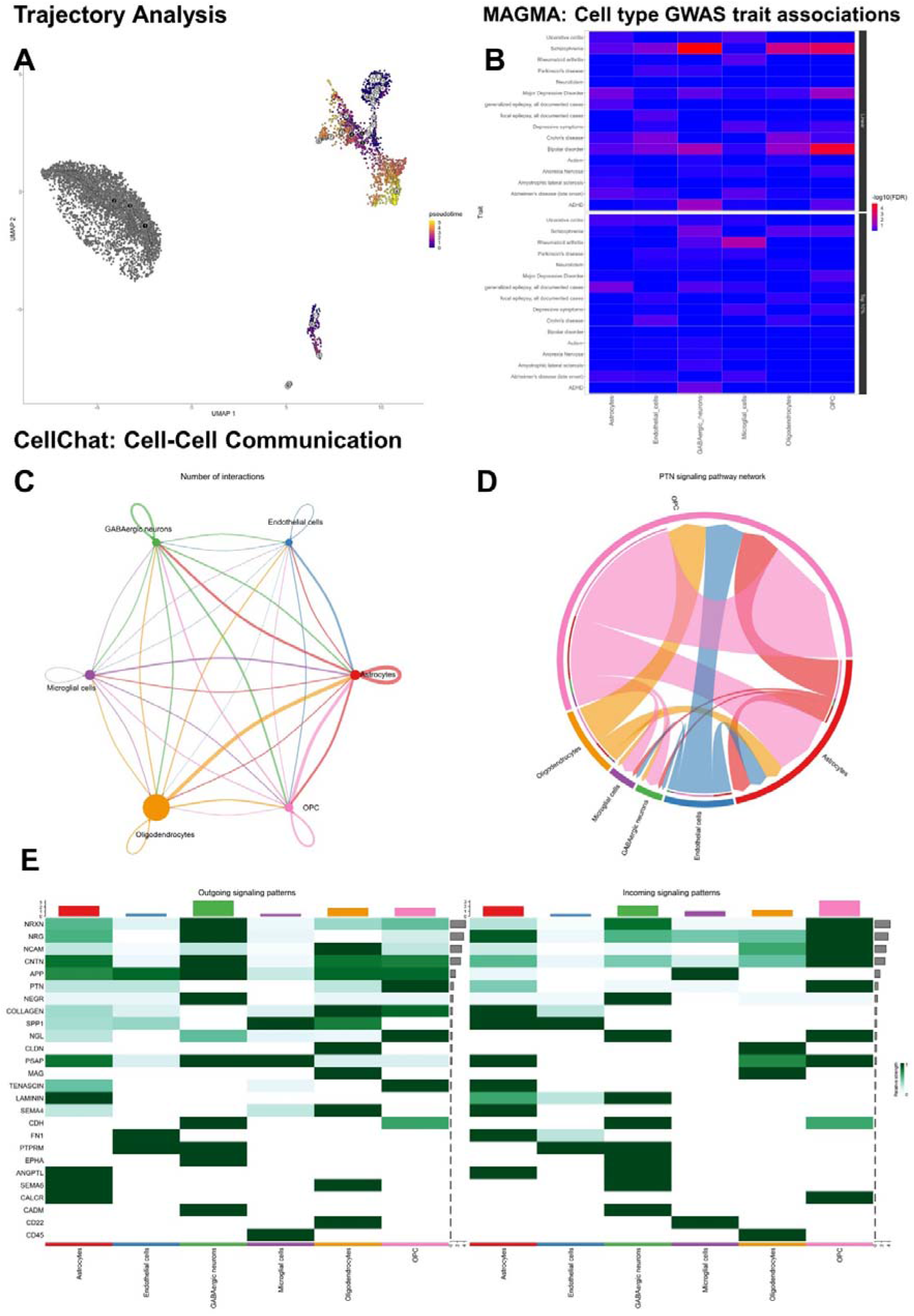
Additional gene network analysis tools available in ICARUS v2.0. Single nuclei from the substantia nigra of 5 human individuals with an absence of neurological disease were analysed with ICARUS v2.0. (A) Trajectory analysis performed with Monocle3 used the oligodendrocyte precursor cells as the root cells (beginning state). Trajectory showcased a gradual pseudotime transition from OPC to astrocytes and microglia. (B) MAGMA cell type GWAS trait associations against several traits spanning immune disease, neurological disorders and psychiatric disease showed an association between GABAergic neurons, OPC and oligodendrocytes with Schizophrenia and bipolar disorder. (C) CellChat cell-cell communication through ligand-receptor pairs showed extensive communication between OPC and astrocytes. Incoming and outgoing signalling pathways are shown as a heatmap (E) with the PTN signalling pathway detailed in the chord plot (D).

To identify cell populations associated with GWAS traits, a MAGMA analysis against several traits spanning immune disease, neurological disorders and psychiatric disease was undertaken (list of GWAS datasets used in analysis listed in Table 2). We have confirmed the association between GABAergic neurons, oligodendrocytes and OPC with the genetic risk of Schizophrenia (P_GABAergic_ = 1.34×10^−5^, P_Oligodendrocytes_ = 2.90×10^−3^, P_OPC_ = 3.40×10^−4^, FDR adjusted) and bipolar disorder (P_GABAergic_ = 0.035, P_Oligodendrocytes_ = 0.071, P_OPC_ = 4.77×10^−5^, FDR adjusted) (Figure 2B); first described by Webber and colleagues (37). Finally, an examination of potential drug-gene interactions, we observe a potential effect for Phorbol 12-myristate 13-acetate (PMA) against the LHFPL3-PTPRZ1-TNR gene cluster described earlier (39,40).

**Table 2.**
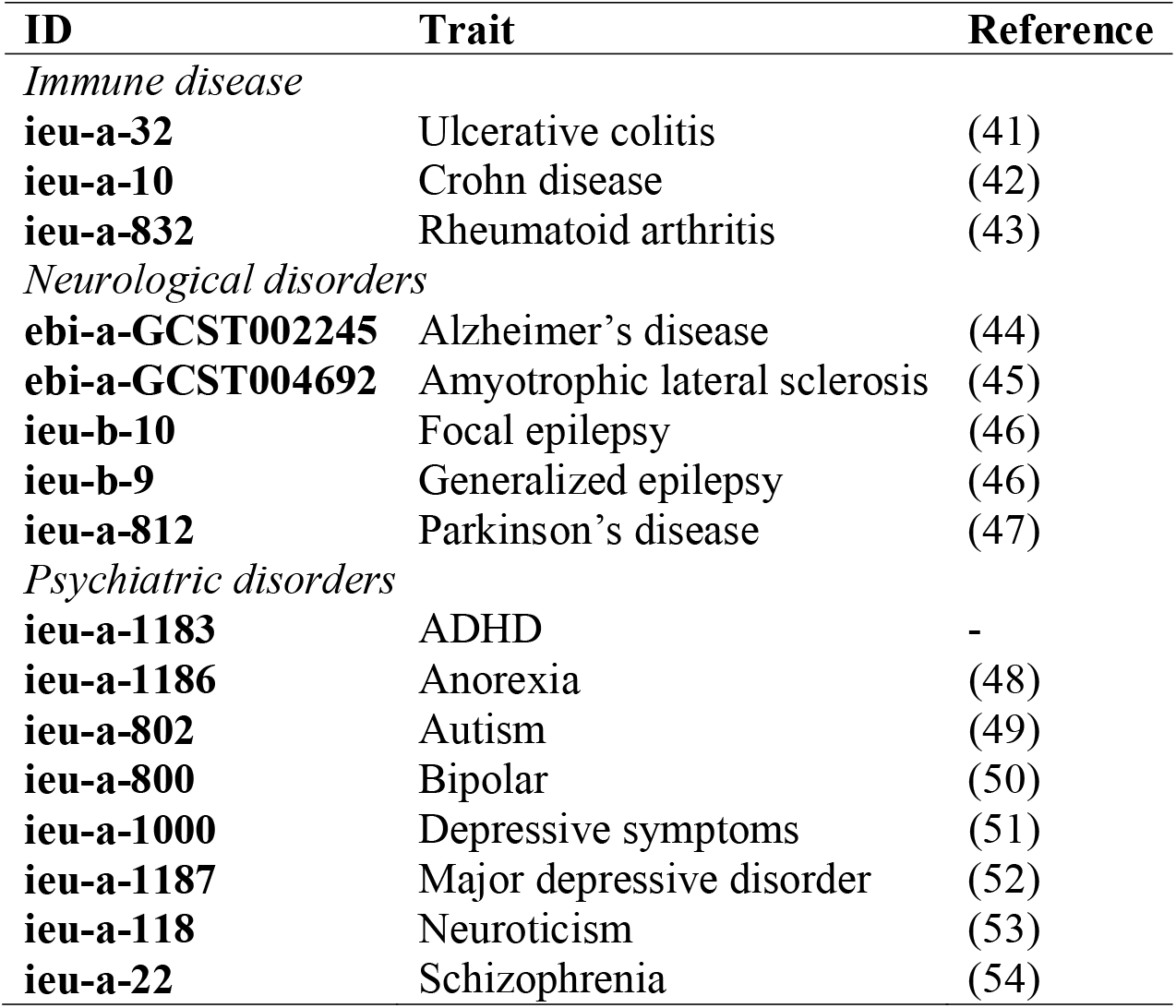
List of GWAS datasets used for MAGMA analysis.

| **ID** | **Trait** | **Reference** |
| --- | --- | --- |
| *Immune disease* |  |  |
| **ieu-a-32** | Ulcerative colitis | (41) |
| **ieu-a-10** | Crohn disease | (42) |
| **ieu-a-832** | Rheumatoid arthritis | (43) |
| *Neurological disorders* |  |  |
| **ebi-a-GCST002245** | Alzheimer's disease | (44) |
| **ebi-a-GCST004692** | Amyotrophic lateral sclerosis | (45) |
| **ieu-b-10** | Focal epilepsy | (46) |
| **ieu-b-9** | Generalized epilepsy | (46) |
| **ieu-a-812** | Parkinson's disease | (47) |
| *Psychiatric disorders* |  |  |
| **ieu-a-1183** | ADHD | - |
| **ieu-a-1186** | Anorexia | (48) |
| **ieu-a-802** | Autism | (49) |
| **ieu-a-800** | Bipolar | (50) |
| **ieu-a-1000** | Depressive symptoms | (51) |
| **ieu-a-1187** | Major depressive disorder | (52) |
| **ieu-a-118** | Neuroticism | (53) |
| **ieu-a-22** | Schizophrenia | (54) |

## Discussion

One of the main reasons for single-cell RNA-seq is to aid in the generation of hypotheses that can be further validated by other experimental techniques. This update to our web server, ICARUS introduces extended tools for data interpretation with a focus on hypothesis generation. ICARUS v2.0 enables users to examine complex gene networks at the single cell level utilising well-established methodologies. We have demonstrated the capabilities of ICARUS v2.0 on a publicly available dataset of the substantia nigra of 5 human individuals with absence of neurological disease (37). Analysis of the oligodendrocyte precursor cell cluster demonstrated the cellular pathway control of cell differentiation and synapse organisation that is likely centred around the co-expression of LHFPL3-PTPRZ1-TNR gene cluster. A clearly defined trajectory from OPCs to astrocytes and microglia mediated by FOS, JUN and JUNB transcription factors was observed that is further supported by increased number of cell-cell signalling from OPCs to astrocytes. Signalling pathways involved include NGL, CNTN and PTN. These observations further contribute to our understanding of the oligodendrocyte progenitor cell lineage and demonstrate the utility of this analysis (55).

Another source of hypothesis generation can be achieved through investigation of potential GWAS trait associated cell populations. Using this dataset of human substantia nigra, we confirm the observation first described by Webber and colleagues (37) of a cell type association between oligodendrocytes, OPCs and GABAergic neuronal gene expression with genetic risk of Schizophrenia and bipolar disorder using the MAGMA methodology. This potentially points to a risk susceptibility for certain cell types underlying these diseases. Finally, the identification of drug-gene targets can facilitate targeted perturbations of key molecular pathways. In the context of dysregulation or disease, this could provide an avenue towards a therapeutic opportunity through repurposed drugs. We identified a possible drug target for the LHFPL3-PTPRZ1-TNR gene cluster expressed in OPCs with a signal transduction activator, Phorbol 12-myristate 13-acetate (39,40).

The updates to ICARUS v2.0 described in this manuscript will provide users with unparalleled resolution into gene networks and understanding of gene regulation in relation to biological traits. The single cell RNA-seq research field is one of the fastest growing fields with over 4,000 research articles published in 2021 alone (Scopus database). ICARUS will continue to see updates as new methodologies are developed to provide the user with a state-of-the-art resource for novel discoveries.

## Data availability

The functionality of ICARUS was demonstrated on a single nuclei RNA-seq dataset of the human substantia nigra from 5 individuals with absence of neurological disease. Data is available from human cell atlas (https://data.humancellatlas.org/explore/projects/996120f9-e84f-409f-a01e-732ab58ca8b9).

## Code availability

ICARUS is available at https://launch.icarus-scrnaseq.cloud.edu.au/. The application is free and open to all users with no login requirement.

R source code of the ICARUS v2.0 shiny app is available at https://github.com/Enjewl/ICARUS. Alternatively, a docker version is accessible through the Docker Hub under the name ‘icarusscrnaseq/icarus’.

## Acknowledgements

We thank the New Zealand–China Non Communicable Diseases Research Collaboration Centre (NCD CRCC). This web server was supported by the Australian National eResearch Collaboration Tools and Resources (Nectar Research Cloud) initiative.

## Contributions

A.J., L.Y., K.L., and R.S., considered the study. A.J., developed the method, implemented the webserver tool, and analysed the data. A.J., wrote the manuscript. A.J., L.Y., K.L., and R.S., supervised the study, tested the webserver and revised the manuscript. All the authors read and approved the final manuscript.

## References

1. Wagner, A., Regev, A. and Yosef, N. (2016) Revealing the vectors of cellular identity with single-cell genomics. Nat Biotechnol, 34, 1145–1160.

2. Jeong, H., Mason, S.P., Barabasi, A.L. and Oltvai, Z.N. (2001) Lethality and centrality in protein networks. Nature, 411, 41–42.

3. Barabasi, A.L. and Oltvai, Z.N. (2004) Network biology: understanding the cell’s functional organization. Nat Rev Genet, 5, 101–113.

4. Jiang, A., Lehnert, K., You, L. and Snell, R.G. (2022) ICARUS, an interactive web server for single cell RNA-seq analysis. Nucleic Acids Research.

5. Song, W.M. and Zhang, B. (2015) Multiscale Embedded Gene Co-expression Network Analysis. PLoS Comput Biol, 11, e1004574.

6. Aibar, S., Gonzalez-Blas, C.B., Moerman, T., Huynh-Thu, V.A., Imrichova, H., Hulselmans, G., Rambow, F., Marine, J.C., Geurts, P., Aerts, J. et al. (2017) SCENIC: single-cell regulatory network inference and clustering. Nat Methods, 14, 1083–1086.

7. Qiu, X., Mao, Q., Tang, Y., Wang, L., Chawla, R., Pliner, H.A. and Trapnell, C. (2017) Reversed graph embedding resolves complex single-cell trajectories. Nature Methods, 14, 979–982.

8. Cao, J., Spielmann, M., Qiu, X., Huang, X., Ibrahim, D.M., Hill, A.J., Zhang, F., Mundlos, S., Christiansen, L., Steemers, F.J. et al. (2019) The single-cell transcriptional landscape of mammalian organogenesis. Nature, 566, 496–502.

9. Trapnell, C., Cacchiarelli, D., Grimsby, J., Pokharel, P., Li, S., Morse, M., Lennon, N.J., Livak, K.J., Mikkelsen, T.S. and Rinn, J.L. (2014) The dynamics and regulators of cell fate decisions are revealed by pseudotemporal ordering of single cells. Nature Biotechnology, 32, 381–386.

10. Jin, S., Guerrero-Juarez, C.F., Zhang, L., Chang, I., Ramos, R., Kuan, C.H., Myung, P., Plikus, M.V. and Nie, Q. (2021) Inference and analysis of cell-cell communication using CellChat. Nat Commun, 12, 1088.

11. de Leeuw, C.A., Mooij, J.M., Heskes, T. and Posthuma, D. (2015) MAGMA: Generalized Gene-Set Analysis of GWAS Data. PLOS Computational Biology, 11, e1004219.

12. Skene, N.G., Bryois, J., Bakken, T.E., Breen, G., Crowley, J.J., Gaspar, H.A., Giusti-Rodriguez, P., Hodge, R.D., Miller, J.A., Muñoz-Manchado, A.B. et al. (2018) Genetic identification of brain cell types underlying schizophrenia. Nature Genetics, 50, 825–833.

13. Freshour, S.L., Kiwala, S., Cotto, K.C., Coffman, A.C., McMichael, J.F., Song, J.J., Griffith, M., Griffith, O.L. and Wagner, A.H. (2021) Integration of the Drug-Gene Interaction Database (DGIdb 4.0) with open crowdsource efforts. Nucleic Acids Res, 49, D1144–D1151.

14. McGinnis, C.S., Murrow, L.M. and Gartner, Z.J. (2019) DoubletFinder: Doublet Detection in Single-Cell RNA Sequencing Data Using Artificial Nearest Neighbors. Cell Systems, 8, 329–337.e324.

15. Hafemeister, C. and Satija, R. (2019) Normalization and variance stabilization of single-cell RNA-seq data using regularized negative binomial regression. Genome Biol, 20, 296.

16. Choudhary, S. and Satija, R. (2022) Comparison and evaluation of statistical error models for scRNA-seq. Genome Biology, 23, 27.

17. Ianevski, A., Giri, A.K. and Aittokallio, T. (2022) Fully-automated and ultra-fast celltype identification using specific marker combinations from single-cell transcriptomic data. Nature Communications, 13, 1246.

18. Zhang, X., Lan, Y., Xu, J., Quan, F., Zhao, E., Deng, C., Luo, T., Xu, L., Liao, G., Yan, M. et al. (2019) CellMarker: a manually curated resource of cell markers in human and mouse. Nucleic Acids Res, 47, D721–D728.

19. Franzén, O., Gan, L.-M. and Björkegren, J.L.M. (2019) PanglaoDB: a web server for exploration of mouse and human single-cell RNA sequencing data. Database, 2019.

20. Hao, Y., Hao, S., Andersen-Nissen, E., Mauck, W.M., 3rd, Zheng, S., Butler, A., Lee, M.J., Wilk, A.J., Darby, C., Zager, M. et al. (2021) Integrated analysis of multimodal single-cell data. Cell, 184, 3573–3587 e3529.

21. Butler, A., Hoffman, P., Smibert, P., Papalexi, E. and Satija, R. (2018) Integrating single-cell transcriptomic data across different conditions, technologies, and species. Nat Biotechnol, 36, 411–420.

22. Satija, R., Farrell, J.A., Gennert, D., Schier, A.F. and Regev, A. (2015) Spatial reconstruction of single-cell gene expression data. Nature biotechnology, 33, 495–502.

23. Stuart, T., Butler, A., Hoffman, P., Hafemeister, C., Papalexi, E., Mauck, W.M., 3rd, Hao, Y., Stoeckius, M., Smibert, P. and Satija, R. (2019) Comprehensive Integration of Single-Cell Data. Cell, 177, 1888–1902 e1821.

24. Korsunsky, I., Millard, N., Fan, J., Slowikowski, K., Zhang, F., Wei, K., Baglaenko, Y., Brenner, M., Loh, P.R. and Raychaudhuri, S. (2019) Fast, sensitive and accurate integration of single-cell data with Harmony. Nature methods, 16, 1289–1296.

25. Aran, D., Looney, A.P., Liu, L., Wu, E., Fong, V., Hsu, A., Chak, S., Naikawadi, R.P., Wolters, P.J., Abate, A.R. et al. (2019) Reference-based analysis of lung single-cell sequencing reveals a transitional profibrotic macrophage. Nature Immunology, 20, 163–172.

26. Darmanis, S., Sloan, S.A., Zhang, Y., Enge, M., Caneda, C., Shuer, L.M., Gephart, M. G.H., Barres, B.A. and Quake, S.R. (2015) A survey of human brain transcriptome diversity at the single cell level. Proceedings of the National Academy of Sciences, 112, 7285–7290.

27. Zhong, S., Zhang, S., Fan, X., Wu, Q., Yan, L., Dong, J., Zhang, H., Li, L., Sun, L., Pan, N. et al. (2018) A single-cell RNA-seq survey of the developmental landscape of the human prefrontal cortex. Nature, 555, 524–528.

28. Baron, M., Veres, A., Wolock, Samuel L., Faust, Aubrey L., Gaujoux, R., Vetere, A., Ryu, Jennifer H., Wagner, Bridget K., Shen-Orr, Shai S., Klein, Allon M. et al. (2016) A Single-Cell Transcriptomic Map of the Human and Mouse Pancreas Reveals Inter- and Intra-cell Population Structure. Cell Systems, 3, 346–360.e344.

29. Lawlor, N., George, J., Bolisetty, M., Kursawe, R., Sun, L., Sivakamasundari, V., Kycia, I., Robson, P. and Stitzel, M.L. (2017) Single-cell transcriptomes identify human islet cell signatures and reveal cell-type-specific expression changes in type 2 diabetes. Genome Res, 27, 208–222.

30. Muraro, Mauro J., Dharmadhikari, G., Grün, D., Groen, N., Dielen, T., Jansen, E., van Gurp, L., Engelse, Marten A., Carlotti, F., de Koning, Eelco J.P. et al. (2016) A Single-Cell Transcriptome Atlas of the Human Pancreas. Cell Systems, 3, 385–394.e383.

31. Segerstolpe, Å., Palasantza, A., Eliasson, P., Andersson, E.-M., Andréasson, A.-C., Sun, X., Picelli, S., Sabirsh, A., Clausen, M., Bjursell, M.K. et al. (2016) Single-Cell Transcriptome Profiling of Human Pancreatic Islets in Health and Type 2 Diabetes. Cell Metabolism, 24, 593–607.

32. He, S., Wang, L.-H., Liu, Y., Li, Y.-Q., Chen, H.-T., Xu, J.-H., Peng, W., Lin, G.-W., Wei, P.-P., Li, B. et al. (2020) Single-cell transcriptome profiling of an adult human cell atlas of 15 major organs. Genome Biology, 21, 294.

33. Wagner, A.H., Coffman, A.C., Ainscough, B.J., Spies, N.C., Skidmore, Z.L., Campbell, K.M., Krysiak, K., Pan, D., McMichael, J.F., Eldred, J.M. et al. (2016) DGIdb 2.0: mining clinically relevant drug-gene interactions. Nucleic Acids Res, 44, D1036–1044.

34. Griffith, M., Griffith, O.L., Coffman, A.C., Weible, J.V., McMichael, J.F., Spies, N. C., Koval, J., Das, I., Callaway, M.B., Eldred, J.M. et al. (2013) DGIdb: mining the druggable genome. Nat Methods, 10, 1209–1210.

35. Russ, A.P. and Lampel, S. (2005) The druggable genome: an update. Drug Discov Today, 10, 1607–1610.

36. Hopkins, A.L. and Groom, C.R. (2002) The druggable genome. Nat Rev Drug Discov, 1, 727–730.

37. Agarwal, D., Sandor, C., Volpato, V., Caffrey, T.M., Monzón-Sandoval, J., Bowden, R., Alegre-Abarrategui, J., Wade-Martins, R. and Webber, C. (2020) A single-cell atlas of the human substantia nigra reveals cell-specific pathways associated with neurological disorders. Nature Communications, 11, 4183.

38. Korsunsky, I., Millard, N., Fan, J., Slowikowski, K., Zhang, F., Wei, K., Baglaenko, Y., Brenner, M., Loh, P.-r. and Raychaudhuri, S. (2019) Fast, sensitive and accurate integration of single-cell data with Harmony. Nature Methods, 16, 1289–1296.

39. Padilla, P.I., Wada, A., Yahiro, K., Kimura, M., Niidome, T., Aoyagi, H., Kumatori, A., Anami, M., Hayashi, T., Fujisawa, J. et al. (2000) Morphologic differentiation of HL-60 cells is associated with appearance of RPTPbeta and induction of Helicobacter pylori VacA sensitivity. J Biol Chem, 275, 15200–15206.

40. Traore, K., Trush, M.A., George, M., Spannhake, E.W., Anderson, W. and Asseffa, A. (2005) Signal transduction of phorbol 12-myristate 13-acetate (PMA)-induced growth inhibition of human monocytic leukemia THP-1 cells is reactive oxygen dependent. Leukemia Research, 29, 863–879.

41. Liu, J.Z., van Sommeren, S., Huang, H., Ng, S.C., Alberts, R., Takahashi, A., Ripke, S., Lee, J.C., Jostins, L., Shah, T. et al. (2015) Association analyses identify 38 susceptibility loci for inflammatory bowel disease and highlight shared genetic risk across populations. Nat Genet, 47, 979–986.

42. Jostins, L., Ripke, S., Weersma, R.K., Duerr, R.H., McGovern, D.P., Hui, K.Y., Lee, J.C., Schumm, L.P., Sharma, Y., Anderson, C.A. et al. (2012) Host-microbe interactions have shaped the genetic architecture of inflammatory bowel disease. Nature, 491, 119–124.

43. Okada, Y., Wu, D., Trynka, G., Raj, T., Terao, C., Ikari, K., Kochi, Y., Ohmura, K., Suzuki, A., Yoshida, S. et al. (2014) Genetics of rheumatoid arthritis contributes to biology and drug discovery. Nature, 506, 376–381.

44. Lambert, J.C., Ibrahim-Verbaas, C.A., Harold, D., Naj, A.C., Sims, R., Bellenguez, C., DeStafano, A.L., Bis, J.C., Beecham, G.W., Grenier-Boley, B. et al. (2013) Metaanalysis of 74,046 individuals identifies 11 new susceptibility loci for Alzheimer’s disease. Nat Genet, 45, 1452–1458.

45. van Rheenen, W., Shatunov, A., Dekker, A.M., McLaughlin, R.L., Diekstra, F.P., Pulit, S.L., van der Spek, R.A., Vosa, U., de Jong, S., Robinson, M.R. et al. (2016) Genome-wide association analyses identify new risk variants and the genetic architecture of amyotrophic lateral sclerosis. Nat Genet, 48, 1043–1048.

46. International League Against Epilepsy Consortium on Complex, E. (2018) Genomewide mega-analysis identifies 16 loci and highlights diverse biological mechanisms in the common epilepsies. Nat Commun, 9, 5269.

47. Simon-Sanchez, J., Schulte, C., Bras, J.M., Sharma, M., Gibbs, J.R., Berg, D., Paisan-Ruiz, C., Lichtner, P., Scholz, S.W., Hernandez, D.G. et al. (2009) Genome-wide association study reveals genetic risk underlying Parkinson’s disease. Nat Genet, 41, 1308–1312.

48. Duncan, L., Yilmaz, Z., Gaspar, H., Walters, R., Goldstein, J., Anttila, V., Bulik-Sullivan, B., Ripke, S., Eating Disorders Working Group of the Psychiatric Genomics, C., Thornton, L. et al. (2017) Significant Locus and Metabolic Genetic Correlations Revealed in Genome-Wide Association Study of Anorexia Nervosa. Am J Psychiatry, 174, 850–858.

49. Cross-Disorder Group of the Psychiatric Genomics, C. (2013) Identification of risk loci with shared effects on five major psychiatric disorders: a genome-wide analysis. Lancet, 381, 1371–1379.

50. Psychiatric, G.C.B.D.W.G. (2011) Large-scale genome-wide association analysis of bipolar disorder identifies a new susceptibility locus near ODZ4. Nat Genet, 43, 977–983.

51. Okbay, A., Baselmans, B.M., De Neve, J.E., Turley, P., Nivard, M.G., Fontana, M.A., Meddens, S.F., Linner, R.K., Rietveld, C.A., Derringer, J. et al. (2016) Genetic variants associated with subjective well-being, depressive symptoms, and neuroticism identified through genome-wide analyses. Nat Genet, 48, 624–633.

52. Wray, N.R., Ripke, S., Mattheisen, M., Trzaskowski, M., Byrne, E.M., Abdellaoui, A., Adams, M.J., Agerbo, E., Air, T.M., Andlauer, T.M.F. et al. (2018) Genome-wide association analyses identify 44 risk variants and refine the genetic architecture of major depression. Nat Genet, 50, 668–681.

53. Genetics of Personality, C., de Moor, M.H., van den Berg, S.M., Verweij, K.J., Krueger, R.F., Luciano, M., Arias Vasquez, A., Matteson, L.K., Derringer, J., Esko, T. et al. (2015) Meta-analysis of Genome-wide Association Studies for Neuroticism, and the Polygenic Association With Major Depressive Disorder. JAMA Psychiatry, 72, 642–650.

54. Schizophrenia Working Group of the Psychiatric Genomics, C. (2014) Biological insights from 108 schizophrenia-associated genetic loci. Nature, 511, 421–427.

55. Zhou, B., Zhu, Z., Ransom, B.R. and Tong, X. (2021) Oligodendrocyte lineage cells and depression. Molecular Psychiatry, 26, 103–117.

