## Supplementary Figure 1 for "Delineation of complex gene expression patterns in single cell RNA-seq data with ICARUS v2.0"

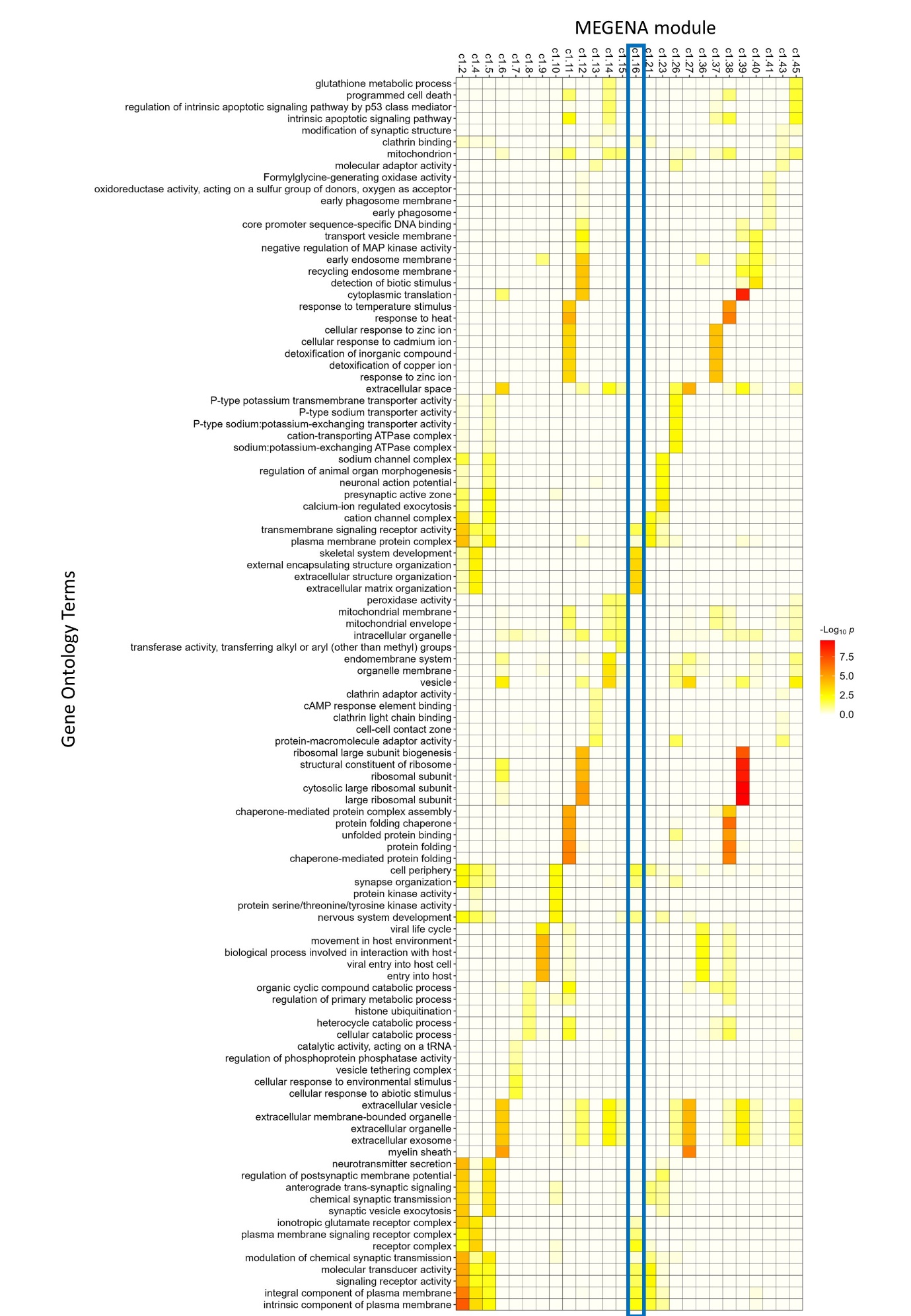

**Supplementary 1** Gene Ontology terms for MEGENA computed gene modules. Module c1_16 (highlighted in blue) involved in extracellular organisation, synapse signalling and developmental morphogenesis was predominately expressed in oligodendrocyte precursor cells.
